# A quantum walk model of Bicoid morphogen formation and interpretation

**DOI:** 10.1101/2023.03.01.530663

**Authors:** Irfan Lone, Carl O. Trindle

## Abstract

The establishment and interpretation of the concentration distribution of the morphogen Bicoid (BCD) is considered crucial for the successful embryonic development of fruit flies. However, the biophysical mechanisms behind the timely formation and subsequent interpretation of the BCD morphogen by its target genes are not yet completely understood. Here a discrete-time, one-dimensional quantum walk model of BCD gradient formation is used to explain both the observed values of diffusivity and its precise interpretation. It is shown that the decoding of positional information from the BCD morphogen by its primary target gene hb, with the observed precision of ∼ 10%, takes a time period of less than a second, as expected on the basis of recent experimental observations. From this the on-rate (*k*_*on*_) for the binding of BCD to its target loci is obtained. Furthermore, the model is also used to explain certain key observations of recent optogenetic experiments concerning the time windows for BCD interpretation. Finally, it is argued that the presented model represents a significant step in the utilization of quantum computation-based techniques in studying the dynamics of biological systems in general and in the field of developmental biophysics in particular.

## I. INTRODUCTION

The establishment of concentration distributions of certain signalling molecules, commonly called morphogens, plays a key role during embryonic development of multi-cellular organisms [1–4]. For example, the concentration distribution of BCD established during the early embryonic development of fruit flies provides the necessary positional information for the proper implementation of the body plan of the organism and has been studied in the greatest detail [5–9]. The timely establishment and precise interpretation of BCD morphogen by its primary target gene hb is considered crucial for the successful development of the fly [9]. Extracellular diffusion accompanied by a unimolecular first order degradation is thought to be the dominant mechanism behind the establishment of BCD morphogen distribution in the fruit fly syncytium [9]. Beginning with the classical Markovian model of morphogen gradient formation by Crick [10], the conventional random walk models of BCD formation have thus employed both continuous as well as discrete-state approximations of a first-order Markov process fundamentally based on the following recursion equation,

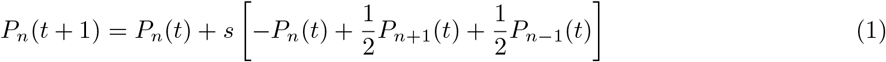

and its continuous limit represented by the diffusion equation [5, 8, 9]. The parameter *s* in such a scheme is a scaling factor that characterizes the probability with which morphogen molecules change their position in time, and we have taken space ∆*x* and time ∆*t* intervals of unit length, a convention which we shall follow throughout [5]. The embryo as a whole, however, is on average a 500 *µ*m long body with the size of each field of cells typically around 100 *µ*m (Fig. 1) [5]. This is clearly a mesoscopic scale [11]. In this regime the system is ideally expected to follow a second-order discrete-state Markovian dynamics, the continuum limit of which is governed by the telegraphist’s equation [12]. The discrete-state biased diffusion approaches [8], which are more in line with this requirement, haven’t, however, yielded a satisfactory theoretical description of the system just like their continuous-state unbiased counterparts [5, 8]. Most importantly, the physical mechanisms behind the precise interpretation of BCD by its primary target gene hb remain still elusive [13–21]. Equilibrium models fail to capture this important aspect of the morphogen in a satisfactory manner and the formulation of increasingly more complex approaches has been suggested [19]. Theories employing detailed balance arguments have also not proven much helpful in this regard [22]. Recent theoretical treatments invoking hypotheses like dynamically changing time windows predict an interpretation time of less than a minute and remain unsubstantiated in vivo [9, 23–25]. However, experimentally BCD is known to bind to chromatin sites in a highly transient manner, with specific binding events lasting on the order of a second, in all portions of the embryo [20]. This demands interpretation times of duration less than a second [20]. Thus, despite extensive numerical and analytical work, the problem of interpretation of the BCD morphogen by its primary target gene hb still has no satisfactory solution, although the formation of its concentration gradient is comparatively somewhat better understood [26].

**FIG. 1.**
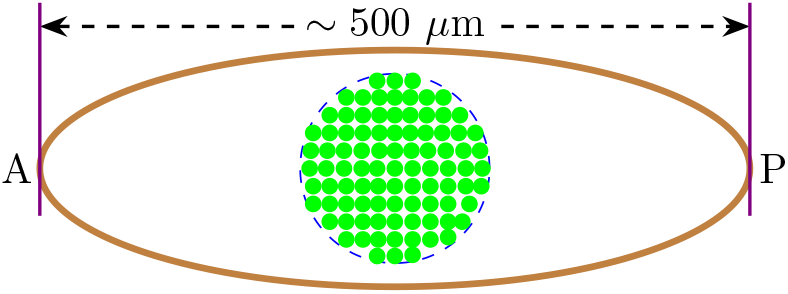
Schematic of the fruit fly embryo showing a typical cell field enclosed in a dashed circle. The diameter of the circle is typically around 100 *µ*m. A and P, respectively, denote the anterior and posterior sides of the syncytium.

In this Letter we present what may be considered as the most straightforward and simplest solution to this perplexing problem on a proof-of-concept basis. We would like to mention at the outset that our model is fundamentally very different from the usual microscopic biased diffusion approaches employing various Markovian kinetic schemes [8] or approximate continuum models derived via rigorous homogenization procedures [27], but its real power lies in that, in addition to bringing out the fundamental limitations of conventional approaches, it can successfully explain some of the most important and elusive features of the BCD-hb system that are currently a matter of intense experimental investigation [19–26]. The treatment to be presented here should, of course, be regarded as a starting point from which to proceed to more rigorous theories. Nonetheless, in this work a conceptual basis is created for describing both the formation as well as interpretation of the BCD morphogen within a novel biophysical framework.

The methods borrowed from quantum mechanics are providing powerful tools for analyzing the dynamics of a wide range of biological phenomena [28–31]. In fact, it has been suggested that biological systems should be described in an infinite dimensional Hilbert space as far as certain self-consistency is required [31]. In the specific case of morphogen gradient formation the problem of time-dependent noise has been analyzed through the use of quantum-mechanical perturbation theory [30]. A Green’s propagator,

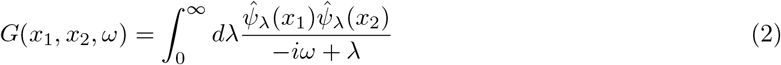

was constructed from the set of eigenfunctions 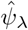 of a linear operator 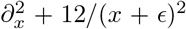 in a non-linear reaction-diffusion model

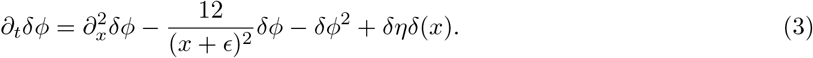

and the iterative solution to the problem, obtained by repeated substitution for the non-linear terms, was then expressed as a sum over tree Feynman diagrams [30]. In Eq. (3) *ϕ*(*x, t*) denotes the local concentartion of morphogen that is produced at the origin at a variable rate *η* and diffuses out to infinity along the positive *x* axis with Fick’s constant *D* accelerating its own degradation at a local rate *fϕ*(*x, t*)^2^. Also *ϵ* = (12*D*^2^*/fη*)^1*/*3^ and *δη*(*t*) is the time-dependent noise that itself is taken to satisfy ⟨*δη*(*t*)⟩ = 0 and ⟨*δη*(*t*)*δη*(*t*′)⟩ = 2*γδ*(*t* − *t*′), the so-called white noise [30], where *δ*(*x*) and *δ*(*t* − *t*′) denote the usual Dirac delta functions.

In view of above, we here develop a quantum walk model of BCD morphogen formation and show that such a model is capable of resolving some of the longstanding and important issues concerning this prototypical morphogenetic system. Specifically, using a modified form of Berg-Purcell limit [32], it is shown that the presented approach can successfully explain the precise interpretation of the BCD morphogen by its target genes and certain key observations of recent optogenetic experiments concerning the time windows for BCD interpretation [33] are also readily explained by the presented model as well.

## II. THEORETICAL METHOD

A special case of the formalism for quantum Markov chains is a framework known as the quantum walk [34]. The quantum walk is a powerful generalization of random walk obtained by imposing quantum mechanics on the walker’s dynamics and works in connection with diffusion in quantum dynamics, in particular, to model the dynamics of a quantum particle on a lattice. It has also garnered numerous applications in diverse areas from understanding the principles underlying various phenomena in physics, chemistry and biology to the designing of search algorithms for quantum computation [35–39]. Here, we develop a quantum walk approach, based on a discrete-time formalism, as a natural framework for incorporating quantum dynamical effects in positional information transfer, as opposed to a classical random walk picture that describes the process in terms of a classical diffusion mechanism. For reasons that will be discussed in the next section, our approach will be purely unitary. However, quantum walks in actual physical systems differ from idealized models of quantum walks in several significant ways. Firstly, Hamiltonians of physical systems typically possess energy mismatches between sites leading to Anderson localization [40]. Secondly, actual quantum walks are subject to relatively high levels of environment-induced noise and decoherence. Therefore, in order to make the development of our approach easier, a basic formalism for a quantum Markov process is briefly reviewed here. A generic representation for the state of a system in quantum mechanics [41] is as follows,

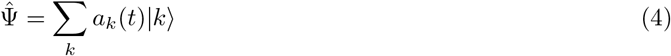

Since, in this work, we will be dealing with discrete times (*t*_*n*+1_ ≥ *t*_*n*−1_ ≥ … ≥ *t*_*n*_ ≥ 0), the time evolution of the probability amplitudes *a*_*k*_ in Eq. (4) is given by the map

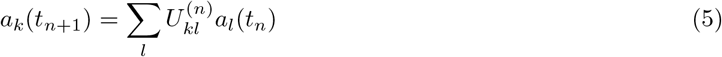

where 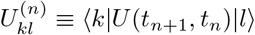 and *U* (*t*_*n*+1_, *t*_*n*_) is the evolution operator connecting the state at time *t*_*n*_ with the one at time *t*_*n*+1_ [42]. In terms of *U*, the occupation probability *P*_*k*_(*t*_*n*_) = |*a*_*k*_(*t*_*n*_)|_2_ can be written as

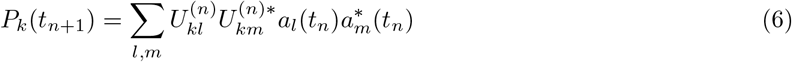

Separating the diagonal terms on the right-hand side, above equation can be written as

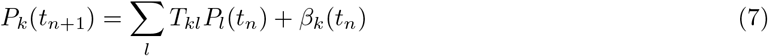

With

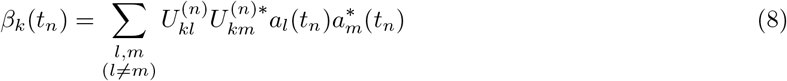

where we have defined 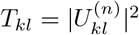 as the transition probability because *T*_*kl*_ ≥ 0 and Σ _*k*_*T*_*kl*_ = Σ _*l*_*T*_*kl*_ = 1.Therefore, if the interference terms (*β*) could be neglected in Eq. (7), the time evolution of the occupation probability would be described by a classical Markovian process in which the transition probability for *k* → *l* in a time interval ∆*t*_*n*_ = *t*_*n*+1_ − *t*_*n*_ is given by *T*_*kl*_. Provided the transition probability is differentiable with respect to time [42], one can then consider the transition probability per unit time *Q*_*kl*_, defined as

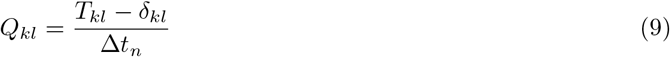

where *δ*_*kl*_ is the Kronecker delta. A straightforward calculation [42] shows that the quantum evolution equation (7) can then be written as

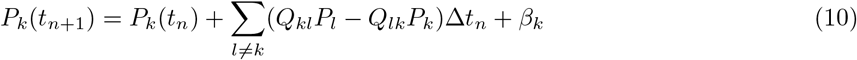

The above equation is composed of two qualitatively different parts: one associated with a Markovian process, i.e., a classical-like diffusion, and the other, *β*_*k*_ associated with quantum interference effects. It is this last term that preserves the unitary character of the evolution. For a purely classical process this term becomes negligible and the evolution is well approximated by

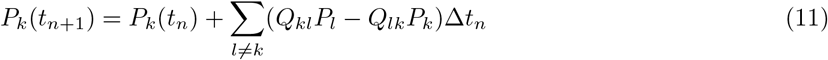

Eq. (11) is sometimes also expressed as

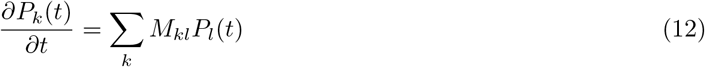

The *M*_*kl*_ denote the Markov transition matrix elements which describe the transition rates between sites *k* and *l* [36, 43]. The form for *U* that we shall employ in our treatment corresponds to a so-called Hadamard walk [44] and is given as

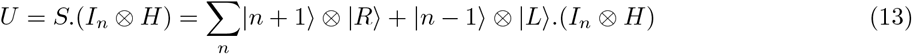

where *H* is the Hadamard operator acting in the coin Hilbert space, *S* a shift operator that acts in the position Hilbert space and conditionally shifts the walker to right or left depending on its chirality state and *I*_*n*_ denotes the identity in the position Hilbert space which itself has a support on the space of eigenfunctions 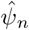 corresponding to the sites *n* on the lattice [44].

### III. RESULTS

#### (A) QUANTUM WALK MODEL OF BCD GRADIENT FORMATION

We begin by considering a hypothetical morphogen composed of particles characterized by an auxiliary chirality degree of freedom that can take two values denoted |*R*⟩ and |*L*⟩, for right and left-handed chirality respectively [44–47]. In order to describe the dynamics of such a particle produced at the origin (*n* = 0) of a one-dimensional infinite lattice (see Fig. 2), all we need is a proper set of recursion equations. Now, unlike a classical walker that occupies a single site at a time, a quantum walker can be in an extended state which is a superposition of all the basis states 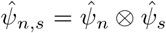. The state of our quantum walker at time *t* is thus represented by

**FIG. 2.**
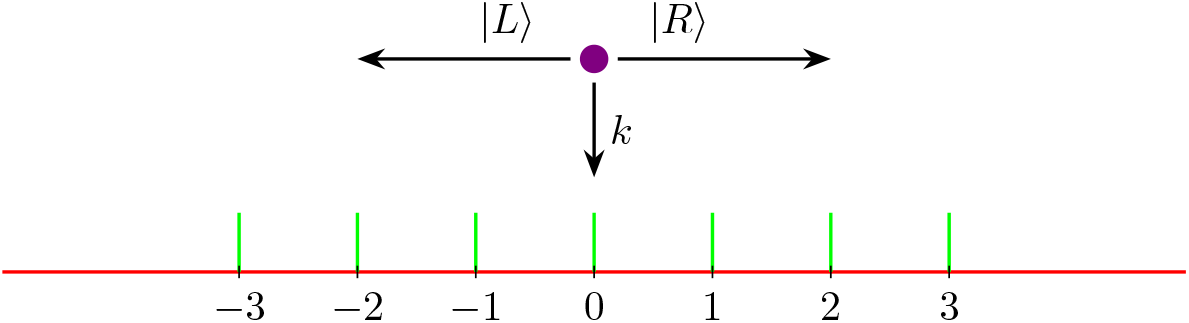
Schematic of the quantum walk model of BCD gradient formation. Each BCD particle is characterized by a chirality degree of freedom that can take two values denoted *R* and *L* for right and left-handed chirality states. A BCD particle of chirality state *R* can move one step to the right at a time while that of chirality *L* can move to the left along the lattice or they can also get degraded, leading to the setting up of a concentration distribution in one-dimensional space. The lattice sites represent the nuclei or positions inside the cytoplasm where there is a finite probability of the morphogen getting degraded.

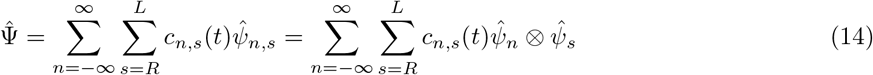

where the amplitudes *c*_*n,s*_ are the complex numbers satisfying Σ_*n,s*_|*c*_*n,s*_|_2_ = 1, which is basically our normalization condition [44]. A commonly employed hypothesis for the establishment of the BCD gradient is that of a steady-state when the rate of synthesis of the morphogen is balanced by its rate of degradation [9]. Under these circumstances there is no net change in the number of morphogen particles and the assumption of normalization of the state of the system is valid. What this means physically is that the net particle number is conserved. The creation and destruction of particles is of no concern once the system reaches a steady-state. Thus, the time dependence of 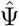 must be such that the normalization is preserved for all time [44]. The probability distribution *P*_*n*_(*t*) is dependent on the chirality state of the walker [47]. Adopting a single-molecule view of the process [8], in which the local concentration of signaling molecules is proportional to a probability *P*_*n*_(*t*) of finding a morphogen molecule at a given site *n* at time *t*, this probability in our case is thus given by

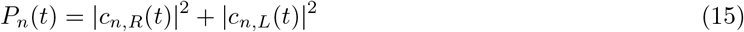

#### (B) QUANTUM NOISE

The formation of the BCD morphogen is fundamentally a noisy process and the degradation of the morphogen is an important contributor to this intrinsic noise in the system [48]. In a classical description [5] this degradation is often modelled as a first-order Markovian death process,

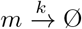

with rate *k*. We shall model it, however, as a special kind of quantum noise [47] that is characterized by a noise rate *p* (see Fig. 3 below). Our quantum noise model implies that this noise rate scales directly with the rate of first order morphogen degradation,

**FIG. 3.**
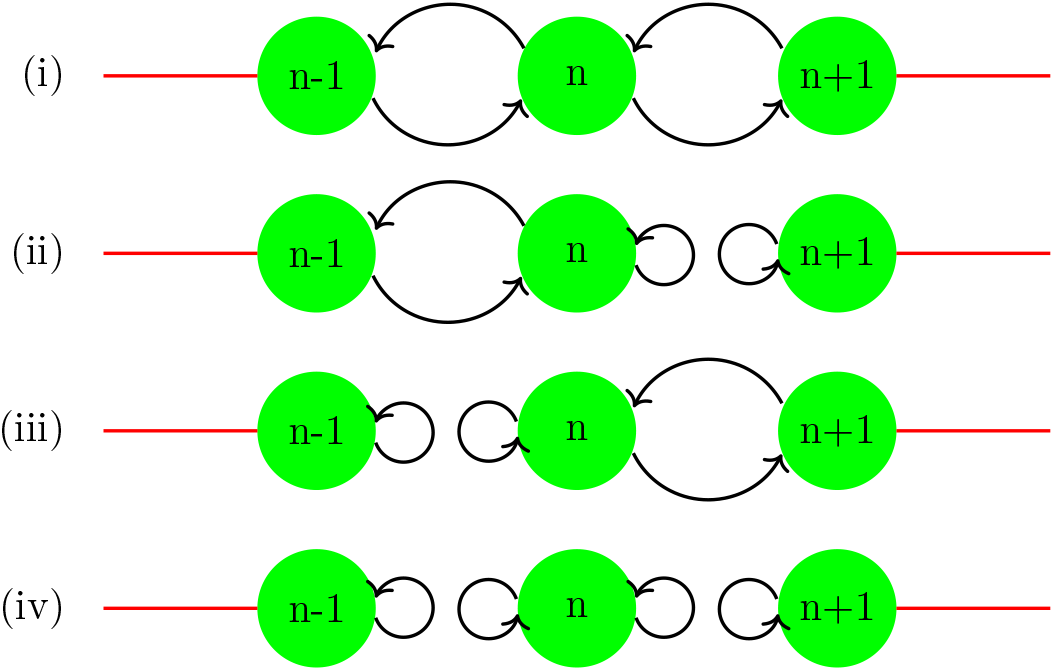
Schematic of quantum noise in the BCD system. The curved arrows depict the flow of probability flux between sites. Lattice sites *n* correspond to nuclei or sites inside the cytoplasm at which the internalization and eventual degradation of BCD molecules can take place. (i) When BCD molecules reside only briefly at a given site *n* they may not get internalized and degraded but instead survive and make a transition to the neighboring sites, *R* to *n* + 1 and *L* to *n* 1. (ii) When a particle of chirality *R* is internalized at site *n* then there is no flow of probability flux towards *n* + 1 and the transition that should have taken place is turned into a self-loop for that step of the walk. However, there is still a flow of probability flux towards *n* − 1 due to the particles of chirality state *L*. (iii) Depicts the reverse of the situation shown in (ii). (iv) When particles of both chirality states reside for appreciably longer periods of time at the site *n* they may both get internalized and eventually degraded. In such a situation there is no flow of probability flux in either direction. In our quantum noise model, we denote the probability per unit time of the particle getting degraded by *p* and the probability of survival by 1 − *p*. So the parameter *p* quantifies noise.

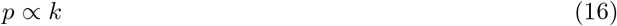

(*p* is inversely related to time and has therefore dimensions of *s*^−1^ which are the dimensions of *k* also [47], therefore, a proportionality between the two is justified in this case.)

#### (C) NON-CLASSICAL MASTER EQUATIONS FOR THE BCD SYSTEM

After taking due recognition of the presence of noise, one can straightforwardly write the following set of four non-classical recursion relations for occupation probabilities for our BCD walker [47] that correspond, respectively, to the situations (i), (ii), (iii) and (iv) shown in Fig. 3,

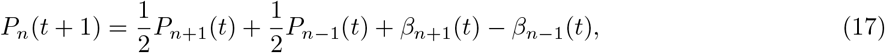

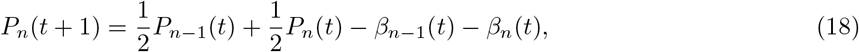

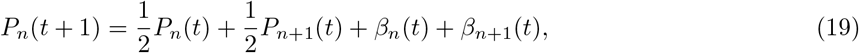

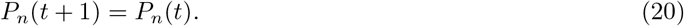

Where 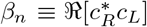 denote the interference terms and the symbol ℜ stands for the real part. If we ignore the *β* terms for the time being and combine the resulting recursion relations into one single equation then a simple scaling argument shows that

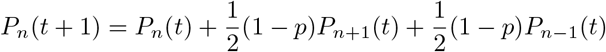

which is the same result as Eq. (1) when *p* = 0. Changing to continuous time and position variables yields the diffusion equation,

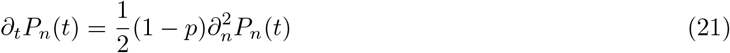

provided we now treat *n* and *t* as continuous variables [46] and define the effective or global diffusivity *D*′ of the system as

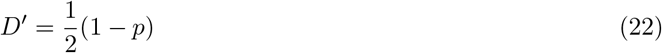

Eq. (21) is the fundamental equation underlying the classical random walk treatments of morphogen gradient formation constructed so far [5, 8, 9]. However, all such models have failed to account for the observed features of the BCD-hb system, for example the time period of interpretation of the morphogen, in an entirely satisfactory manner [13, 21, 23].

#### (D) LOCAL AND GLOBAL DIFFUSIVITY OF BCD

For a purely classical walk of the morphogen *p* = 0 and, as Eq. (22) shows, the effective diffusion coefficient for the system is the same as that for an unbiased classical random walk (∼ 0.5*µm*^2^*s*^−1^) [49]. However, for a non-zero noise rate *p* the value of *D*′ is even below the unbiased random walk value of ∼ 0.5*µm*^2^*s*^−1^. Gregor *et al*. have reported a value of ∼ 0.3*µm*^2^*s*^−1^ for the effective diffusivity of Bcd in nuclear cycle 14 embryos using FRAP as well as certain indirect calculations [50]. This observation is fully consistent with Eq. (22) considering the strong noise levels (*p* ∼ 0.4 from Eq. (22)) that are expected in the system at this stage in embryonic development. Now, in order to account for the high values of local diffusivity observed through FCS measurements during nuclear cycles 12–14 [51], and to explain the observed precision of patterning processes in the system, Eqs. (17)–(20) must be solved fully without neglecting the *β* terms. A numerical solution can be implemented [47] to show that,

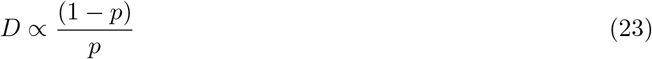

Such a diffusivity is called local diffusivity *D* for the system. As can be clearly seen from Eq. (23), the local diffusivity of BCD system is not constant but varies with the noise in the system reaching values much larger than the classically possible ones when the noise levels are very small. It is also interesting to discuss here the biological implications of such small noise levels. Higher noise levels, due to faster degradation rates, can have the consequence of decreasing the effective mobility of the morphogen such that less and less particles reach a given location in the tissue compromising the consequent transfer of positional information. Thus, it is beneficial for the system to keep the degradation rates very low because increased degradation also increases noise. This can have negative consequences for the ongoing development processes. It is therefore reasonable to suggest that the fly embryo controls complex biochemical and biophysical processes of development by modifying degradation, and hence noise, rates. This also explains the very low degradation rates of the morphogen observed experimentally [48].

Based on the currently available values of diffusivity, which are measured around nuclear cycle 14, BCD is classified as a mid-range morphogen [52]. However, recent work has detected significant concentration of BCD in the posterior-most parts of the embryo [20]. For example, lattice light sheet microscopy studies have revealed highly dynamic BCD molecules in the posterior end of the embryo with a diffusive-like kinetic behavior [20]. With the observed low levels of degradation, and hence noise, that are known in the literature [48, 53, 54] the possible values of diffusivity for BCD as per Eq. (23) can be an order of magnitude higher than are currently known. Such large diffusion coefficients for morphogens are not an uncommon occurrence and values as high as 50 − 80*µm*^2^*s*^−1^ have been measured in different cellular contexts [52]. Lower noise rates, especially in earlier nuclear cycles, will yield very large diffusion coefficients for the system. Most quantitative studies of fruit fly development have, however, focused on nuclear cycle 14 embryos where the nuclei are easily accessible for imaging [9]. A very high diffusion rate predicted by our model and confirmed experimentally can also solve the problem of generating an appropriate gradient sufficiently fast for significantly larger patterning fields and BCD could also become classified as a long-range morphogen like Nodal, Lefty and FGF8 in the zebrafish embryo [52].

Thus, as per above analysis there is no single characteristic value of local diffusivity for the BCD system but a range of possible values and the high values of BCD diffusivity (5 − 10*µm*^2^*s*^−1^) obtained through FCS measurements [51] around nuclear cycles 12–14 are a consequence of Eq. (23). Furthermore, higher and higher values of diffusivity should be expected when the noise levels are very small especially in early nuclear cycles. It is pertinent to mention here that a recent analysis has shown that a temporally varying diffusivity for BCD is better at capturing the dynamics of gradient formation for this morphogen within the SDD scheme [26]. However, the precise mechanism behind such temporal dependence remains unknown [26]. Present analysis suggests an explanation of this observation. Therefore, if the system can vary its diffusivity depending upon its requirement by modulating noise, then there is no single fixed value of local diffusivity for the system but rather a range of possible values. What this means, in other words, is that the BCD protein mobility changes with time and spatial position in the embryo. It is this crucial piece of information contained in Eq. (23) that we use here to account for the observed precision of gene expression patterns initiated by the BCD system. One of the simplest and best studied systems in this regard is the BCD-hb system in the early fruit fly embryo [55].

#### (E) PRECISE INTERPRETATION OF THE BCD MORPHOGEN

Roughly, the timescale of BCD driven pattern formation in the fly embryo can be divided into two sub-intervals. First the 90 minutes prior to nuclear cycle 10 are utilized by the system for the establishment of the BCD distribution in the embryo [50]. From nuclear cycle 10 onward the BCD concentration inside the nuclei remains stable from cycle to cycle [50]. Furthermore, it is now well established that the quantitative relationship between the statistics of hb expression levels is consistent with a model in which the dominant source of noise are the diffusive fluctuations associated with the random arrival of BCD molecules at their target sites [56]. This noise is quantitatively expressed by the Berg-Purcell limit usually given as

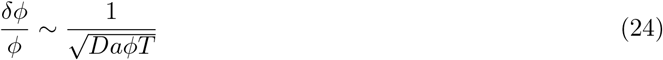

in which *a* denotes the size of the BCD receptor, *ϕ*(*x, t*) is the local concentration of the morphogen, *D* is its diffusivity in the matrix surrounding the receptor and *T* is the duration of signal integration [13]. Intriguingly, what Gregor *et al*. discovered is that the observed precision of ∼ 10 % in the readout of the BCD concentration by the nuclei was inconsistent with the predictions of this fundamental physical limit for diffusive processes [13]. To circumvent this problem, the idea of spatial averaging by the nuclei was introduced in order to increase the precision to the desired level [13]. Over the years the mechanism of Gregor *et al*. has been widely applied [13–19] but the problem of interpretation of BCD gradient by its primary target gene hb still remains poorly understood [19–25]. In fact a recent theoretical analysis has shown that spatial averaging is *a priori* dispensable but the treatment given is unwieldy and introduces another *adhoc* hypothesis of dynamically changing time windows [23]. Most importantly, it predicts an interpretation time much higher than is supported by current experimental observations based on lattice light sheet microscopy studies [20]. Since, as per Eq. (23), the diffusivity of BCD is very large for low noise levels in the system and, as we have seen above, the system could vary its diffusivity depending upon its requirement by modulating noise, it is thus quite conceivable that a significant fraction of BCD molecules could be moving much faster even after nuclear cycle 10 given the very low observed degradation rates (*k*∼ 10^−4^*s*^−1^) of the BCD morphogen in the system [26, 53, 54]. Therefore, if we substitute Eq. (23) in Eq. (24) we obtain

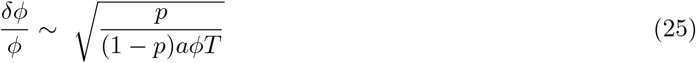

and Eq. (25) is our modified Berg-Purcell limit. We use Eq. (25) to explain how BCD can be precisely decoded by its primary target gene hb as follows. In order to estimate a typical value of noise that is sufficient to achieve the desired level of precision in the system we use a simple order of magnitude calculation based on the observed values of the first order degradation rates in the system. Taking a typical value of the measured BCD degradation rate (*k*∼ 10^−4^*s*^−1^) around nuclear cycles 10–14 [26, 48, 53, 54] and using (6) that *p* ∝ *k*, we can therefore write *p* ∼ *k* ∼ 10^−4^*s*^−1^. Substituting this value of noise and the values of concentration *ϕ* and receptor size *a* from [13] into Eq. (25) for the modified Berg-Purcell limit

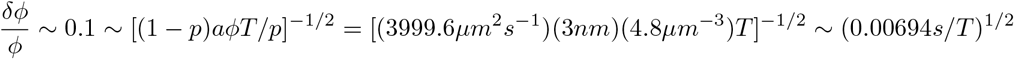

shows that the decoding of positional information from the BCD morphogen by its primary target gene hb, with the observed precision of ∼ 10%, takes less than a second (*T* ∼ 0.7*s*). From this one can readily calculate the on-rate of the BCD morphogen for its target loci as

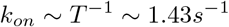

which is a very moderate on-rate. This is significant considering that recent experiments have shown BCD binding to chromatin sites in a highly transient manner, with specific binding events lasting on the order of a second, in all portions of the embryo with a rapid off rate such that its average occupancy at target loci is on-rate dependent [20].

#### (F) TIME WINDOWS FOR BCD INTERPRETATION

By subjecting the fly embryos to illumination at precisely defined stages, distinct time windows for the interpretation of BCD have been defined through optogenetic experiments [9, 33]. It has been demonstrated that temporal integration is indeed a necessary process in BCD decoding with the integration time window covering as long as 90 minutes from nuclear cycle 10 until the end of nuclear cycle 14 [33]. Interestingly, the nuclei exposed to highest local BCD concentration, those on the anterior side, integrated BCD signal for the longest duration while those receiving lower BCD dosage, that is nuclei closer to the posterior side, required BCD inputs for comparatively shorter times in order to arrive at correct cell fates [9, 33]. Now, a spatio-temporally nonuniform degradation of BCD is responsible for a non-uniform distribution of noise in the system. This non-uniformity is due to the higher concentration of BCD on the anterior side of the embryo than that towards the posterior side. This is despite the fact that the net concentration of BCD inside the nuclei at a given position along the AP-axis remains constant within ∼ 10% accuracy from nuclear cycle 10 onward [50]. The net effect of all these increased noise processes on the anterior is to reduce the diffusivity of the system, the (1 − *p*)*/p* term in Eq. (25), more on the anterior side and consequently the nuclei on this side need to integrate the signal for a longer duration (higher *T* values) in order to reach the desired level (∼ 10%) of precision in signal interpretation. Hence the resulting different time windows.

## IV. DISCUSSION

According to Wolpert, the embryo should be computable “if a level of complexity of description of cell behavior can be chosen that is adequate to account for development but that does not require each cell’s detailed behavior to be taken into account”[57]. We view the presented treatment as an important step towards the realisation of this goal. This study is, to the best of our knowledge, the first to employ the formalism of quantum walks in explaining the dynamics of morphogen gradient formation and interpretation. On the other hand, quantum walks are now increasingly being exploited as important tools in diverse areas from understanding the principles underlying various phenomena in physics, chemistry and biology to quantum computation and information processing tasks and have recently been shown to constitute a universal model for computation [35–39]. However, the transmission and processing of information inside biological systems is much faster than is attainable at the present stage of computation [31]. Seen from this fundamental viewpoint, developing the mathematical models of living systems that incorporate quantum computation-based methods is expected to yield a better insight into the dynamical principles governing life.

The above discussion focused on a very simple model of BCD gradient formation and its subsequent interpretation by the nuclei. The environment in which the BCD distribution forms is highly dynamic and heterogeneous. In fact, the Markovian behavior is always an idealization in the description of the dynamics of open systems with non-Markovianity being non-negligible in a number of different scenarios especially biological [58]. However, the system we dealt with in this work is unique and special in so many ways that even a Markovian description suffices. The BCD system is essentially a one-dimensional problem since patterning systems along different axes of the embryo are independent of each other [59]. Hence a simple one-dimensional quantum walk treatment becomes possible. It would be interesting to see whether a similar type of analysis can be carried out for other morphogens, particularly the gradient of the morphogen DPP established orthogonal to BCD along the dorso-ventral axis of the embryo [60].

Despite the apparent simplicity of our approach it makes, in addition to explaining the observed features of the system, certain new and useful predictions. The presented analysis, however, should not be taken as an attempt to invalidate the SDD model completely, which has after all a good deal of experimental evidence in its favor [26]. Rather what our treatment does is explore the dynamics underlying the SDD scheme in more detail. In other words, while as the SDD model provides a low-resolution picture of the patterning system [1] our model represents a high-resolution picture of its underlying dynamics that gives rise to the SDD behavior at larger time and distance scales [5]. Most importantly, as is clear from Eq. (25), we find that the problems of gradient formation and interpretation are interconnected processes, at least in the case of BCD-hb system. In this regard our model is a very rare instance of a long desired approach [62] that incorporates both the dynamics of morphogen gradient formation and its subsequent interpretation. Specifically, we find that the embryo does not need to rely on complex mechanisms like spatial averaging [13, 14] and dynamically changing time windows [23] in order to generate precise boundaries of gene expression in a short span of time. Rather, the very large values of diffusivity attained by the system are sufficient to achieve the desired level of precision in the patterning processes.

## V. CONCLUSION

We developed a one-dimensional discrete-time quantum walk model for understanding the dynamics of BCD gradient formation and interpretation in the early fruit fly embryo. The approach also provided an explanation of recent optogenetic experiments concerning the time windows for BCD interpretation. It allowed us to explain both the observed values of diffusivity and the precise interpretation of the BCD morphogen by its primary target gene hb. It is found that the very large values of diffusivity are responsible for generating precise boundaries of gene expression in a short span of time. This is in contrast to the conventional treatments that require complex mechanisms like spatial averaging and dynamically changing time windows to achieve the desired level of precision in the patterning processes. Finally, we believe that our findings should provide motivation for an exciting new line of experimental inquiry that can yield a better understanding of the factors that play a decisive role in the processes of embryonic development.

## VI. ACKNOWLEDGEMENTS

IL acknowledges the facility provided by CQRC Canada. Correspondence with Anatoly B. Kolomeisky over the past many years regrading various aspects of modelling the BCD morphogen is also acknowledged. Last, but not least, the constructive criticism of certain anonymous reviewers is greatly acknowledged.

